# A single-cell transcriptomic atlas of the pigtail macaque placenta in late gestation

**DOI:** 10.1101/2025.08.15.669966

**Authors:** Amanda Li, Richard Li, Hazel Huang, Hong Zhao, Briana Del Rosario, Miranda Li, Edmunda Li, Andrew Vo, Gygeria Manuel, Orlando Cervantes, Raj P. Kapur, Jeff Munson, Austyn Orvis, Michelle Coleman, Melissa Berg, Britni Curtis, Brenna Menz, Jin Dai, Inah Golez, Solomon Wangari, Chris English, Audrey Baldessari, Lakshmi Rajagopal, John Cornelius, Kristina M. Adams Waldorf

**Affiliations:** Department of Obstetrics and Gynecology, University of Washington, Seattle, Washington, United States of America; Case Western Reserve, Cleveland, Ohio, United States of America; Columbia University, New York, New York, United States of America; School of Medicine, University of Washington, Seattle, Washington, United States of America; Morehouse School of Medicine, Atlanta, Georgia, United States of America; Department of Global Health, University of Washington, Seattle, Washington, United States of America; Department of Pathology, University of Washington, Seattle, Washington, United States of America; Department of Pathology, Seattle Children’s Hospital, Seattle, Washington, United States of America; Department of Psychiatry, University of Washington, Seattle, Washington, United States of America; Center for Global Infectious Disease Research, Seattle Children’s Research Institute, Seattle, Washington, United States of America; Washington National Primate Research Center, Seattle, Washington, United States of America

**Keywords:** Placenta, fetus, pigtail macaque, nonhuman primate, decidua, chorionic villous, chorioamniotic membrane

## Abstract

The placenta is a complex organ with multiple immune and non-immune cell types that promote fetal tolerance and facilitate the transfer of nutrients and oxygen. The nonhuman primate (NHP) is a key experimental model for studying human pregnancy complications, in part due to similarities in placental structure, which makes it essential to understand how single-cell populations compare across the human and NHP maternal-fetal interface. We constructed a single-cell RNA-Seq (scRNA-Seq) atlas of the placenta from the pigtail macaque (*Macaca nemestrina*) in the third trimester, comprising three different tissues at the maternal-fetal interface: the chorionic villi (placental disc), chorioamniotic membranes, and the maternal decidua. Each tissue was separately dissociated into single cells and processed through the 10X Genomics and Seurat pipeline, followed by aggregation, unsupervised clustering, and cluster annotation. Next, we determined the maternal-fetal origins of cell populations and analyzed single-cell RNA trajectory, Gene Ontology enrichment, and cell-cell communication. Single-cell populations in the pigtail macaque were strikingly similar in their identity and frequency to those found in the human placenta, including cells from trophoblast, stromal cell, immune, and macrophage lineages. An advantage of our approach was the deep sequencing of three tissues at the maternal-fetal interface, which yielded a rich diversity of common and rare single-cell populations. The third-trimester pigtail macaque single-cell atlas enables the identification of cellular subclusters analogous to those in humans and provides a powerful resource for understanding experimental perturbations on the NHP placenta.

## Introduction

The placenta is an essential, temporary organ responsible for providing nutrients, oxygen, and a protective immune environment to the fetus without which a mammal could not survive. Placentas vary markedly across species, yet their diverse structures perform the same critical functions to support the developing fetus.(1, 2) Pregnant animal models are powerful tools to investigate the pathogenesis of human reproductive disorders, especially when the placental biology of the model species closely resembles that of humans. Macaques, belonging to the genus *Macaca*, are commonly used as animal models in pregnancy research, largely due to their close similarity with humans in placental structure, fetal development, and physiology of labor onset. Rhesus macaques (*Macaca mulatta*), pigtail macaques (*Macaca nemestrina*), and cynomolgus macaques (*Macaca fascicularis*) are frequently used to study the impact of infectious diseases(3–15) and reproductive disorders(16, 17) on fetal development(18–21), placental health(22–25), and perinatal outcomes(26, 27). Although these species have served as models of the impact of experimental stimuli on human reproduction and pregnancy, the degree to which they have similar populations of placental immune and non-immune cells (i.e., epithelial cells, mesenchymal cells, stromal, and trophoblast cells) is incompletely known.

The overall structure of the macaque placenta is very similar to that in humans consisting of a bi-discoid villous placenta with a tri-hemochorial interface.(28) Occasionally, the macaque placenta may also form a single disc, as in humans. Key differences in placental cellular composition between humans and macaques during the second and third trimesters include shallower trophoblast invasion in macaques (2), a continuous trophoblastic shell in macaques versus a less uniform structure in humans(29, 30), and the rarity in macaques of interstitial extravillous trophoblasts(31), which are common in humans. Despite these differences, the macaque placenta closely resembles the human placenta, sharing similar trophoblast lineages, villous architecture, the production of human chorionic gonadotropin (hCG)-like molecules and progesterone, spiral artery remodeling, and mechanisms that support maternal-fetal immune tolerance. The total gestation time of the macaque pregnancy from the day of conception is approximately 167 days(32), 172 days(33), and 164 days(34) for rhesus, pigtail, and cynomolgus macaques, respectively. In contrast, human gestation is approximately 100 days longer than the macaque pregnancy (266 days from conception or 280 days from the first day of a woman’s last menstrual period).

Single-cell RNA sequencing (scRNA-Seq) has enabled the investigation of individual cell transcriptomes and the analysis of cellular heterogeneity in complex tissues, including the placenta.(35, 36) Several scRNA-Seq atlases of the human placenta have been published, mostly describing cell populations in the first and second trimesters(37–42), but also in the third trimester(43–45). scRNA-Seq datasets from nonhuman primate (NHP) placentas are comparatively rare with a scRNAseq analysis of the cynomolgus macaque placenta at different stages of gestation, scRNA-Seq of CD14+ cells in a ZIKV-infected pregnancy, and a pre-print from rhesus macaque placental organoids.(46–48) As the nonhuman primate is a common model of pregnancy disorders, especially in the third trimester, a normal placental atlas in late gestation and at high resolution is valuable.

The objective of our study was to establish a scRNA-Seq atlas of the pigtail macaque placenta from the late second and third trimesters, with high-resolution characterization of cell populations at the maternal-fetal interface within the maternal decidua, placental villous tissue, and chorioamniotic membranes. We have used the pigtail macaque as an NHP model to investigate the pathogenesis of several infectious diseases in pregnancy and different pathways of preterm labor.(5, 6, 11, 13, 26, 27, 49–53) Establishing the frequency and distribution of immune and non-immune cell populations in the macaque placenta and at the maternal-fetal interface will enable comparison with scRNA-Seq data from other models of infectious diseases or reproductive disorders.

## Results

### Comparative Anatomy of the Pigtail Macaque and Human Placentas

Creation of a scRNA-Seq atlas of the pigtail macaque placenta required an appreciation of the anatomical similarities and differences with human placentas (Fig. 1). Overall, the architecture of the placental chorionic disc is remarkably similar between the two species with the human placenta (Fig. 1E) being much thicker than the macaque (Fig. 1A). At higher magnification, the placental chorioamniotic membranes (Fig. 1B, 1F), chorionic villi (Fig. 1C, 1G), and basal plate (Fig. 1D, 1H) exhibit similar architectural patterns. An important difference between the macaque (Fig. 1D’) and human (Fig. 1H’) placentas is the absence of cytokeratin 7-positive interstitial extravillous trophoblast cells in the macaque basal plate, a finding that we confirmed in our own model and has been previously reported in closely related nonhuman primate species.(54–56) Based on these observations, we did not expect to annotate extravillous trophoblast cells in our scRNA-Seq data.

**Figure 1.**
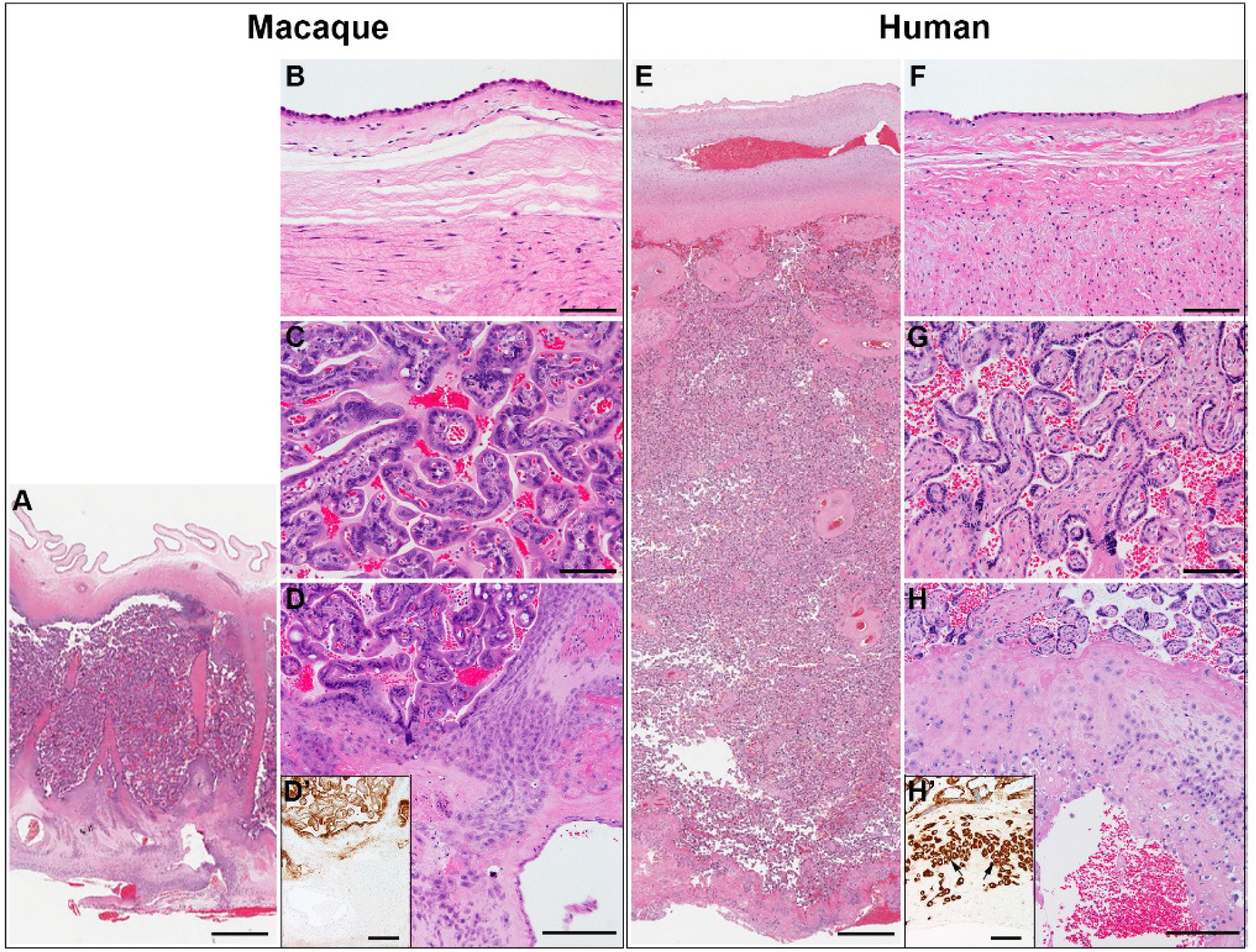
Comparative microscopic anatomy of pigtail macaque and human placentas. Full-thickness sections (fetal surfaces at top) of the chorionic discs from third-trimester pigtail macaque (A-D’) and human (E-H’) placentas are shown with higher magnification images of the chorioamniotic membranes (B, F), chorionic villi (C, G), and the basal plate (D, H). Immunostaining for cytokeratin 7 was performed to identify interstitial extravillous trophoblast cells in the basal plate, which were present in the human (arrows in H’), but not the macaque (Fig. 1D). Scale bars: A, E: 1 mm; all others, 100 µm.

### Study Design and Analysis of the Biological Variability Across Samples

To determine the single-cell RNA transcriptional profiles of the pigtail macaque placenta, we collected scRNA-Seq data separately from chorionic villous tissue, chorioamniotic membranes, and the adjacent maternal decidua. Tissues were obtained from 10 healthy, uninfected pregnant pigtail macaques that had either undergone surgical catheterization and saline inoculation of the amniotic cavity and choriodecidual space (N=5) or media inoculation of the forearm (N=5; Table S1). These animals represented controls for infectious disease studies. All animals were delivered by Cesarean section, and none developed preterm labor or other perinatal complications. Chorionic villous tissue from the placental disc (DISC), chorioamniotic membranes (CAM) and decidual tissue (DEC) were harvested immediately after delivery and processed separately through a scRNA-Seq pipeline to integrate the data, annotate single cell populations, and determine the tissue and maternal or fetal origin for each cluster.

While sequencing freshly dissociated tissue enhanced the quality of our data, it also introduced potential batch effects, as samples were sequenced on separate chips. We explored changes in the Biological Coefficient of Variation (BCV, Fig. 2) of the log gene counts per million (CPM) split across tissue types to determine whether the Harmony software tool adequately handled variance due to multiple sequencing runs. Prior to Harmony integration, the BCV was high (∼2%) with minimal dispersion of the average log CPM (Fig. 2A, 2C, 2E). After Harmony integration, the BCV was approximately 0.5% across the tissues with a greater spread in average log CPM count (Fig. 2B, 2D, 2F). We considered this level of data dispersion to be acceptable, given that each biological sample was sequenced fresh, which led to batch effects.

**Figure 2.**
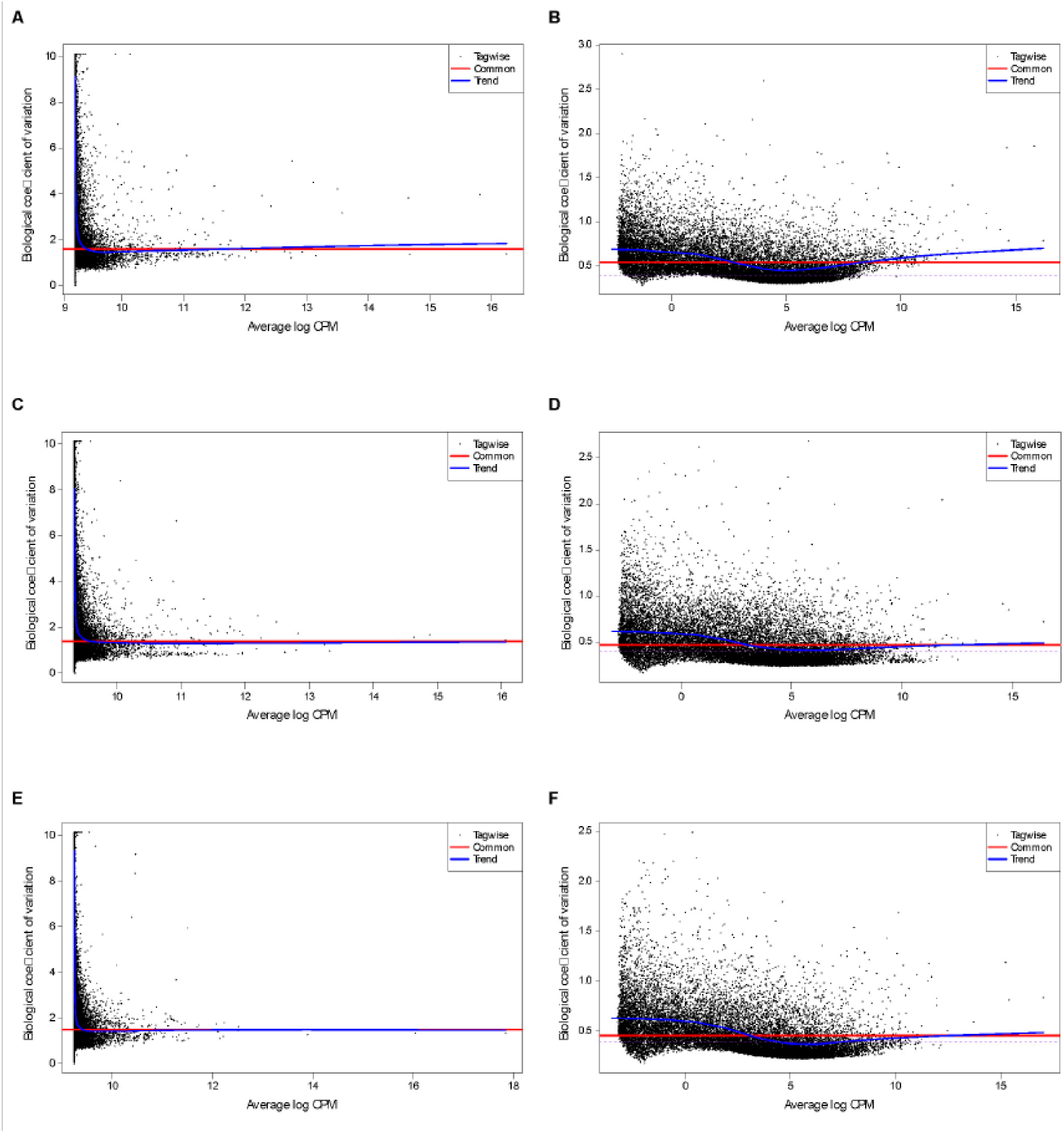
Biological Coefficient of Variation on Pseudobulked Data Before and After Data Harmonization. Plots reflecting the biological coefficient of variation (BCV) are shown using pseudobulked data for DISC, CAM, and DEC, before (A: DISC; C: CAM; E: DEC) and after (B: DISC; D: CAM; F: DEC) data integration and batch correction using Harmony software. The y-axis is the BCV, and x-axis represents average log counts per million (CPM), a normalization method to scale the raw read counts to the number of reads mapping to a gene per million total mapped reads in a sample.

### Clustering, Annotation, and Cell Origin

Unsupervised clustering analysis and manual annotation initially yielded 33 clusters, which included two large groups of immune cells. A second round of unsupervised clustering specifically on these two groups was necessary to achieve greater resolution of the immune cell populations, which could then be easily identified using canonical markers found in the literature or cell marker databases.(57) The additional unsupervised clustering on the immune cell and macrophage groups yielded an additional 30 clusters. Once populations were annotated and similar populations merged, a total of 33 unique cell populations were identified and annotated in a uniform manifold approximation and projection (UMAP) (Fig. 3A).

**Figure 3.**
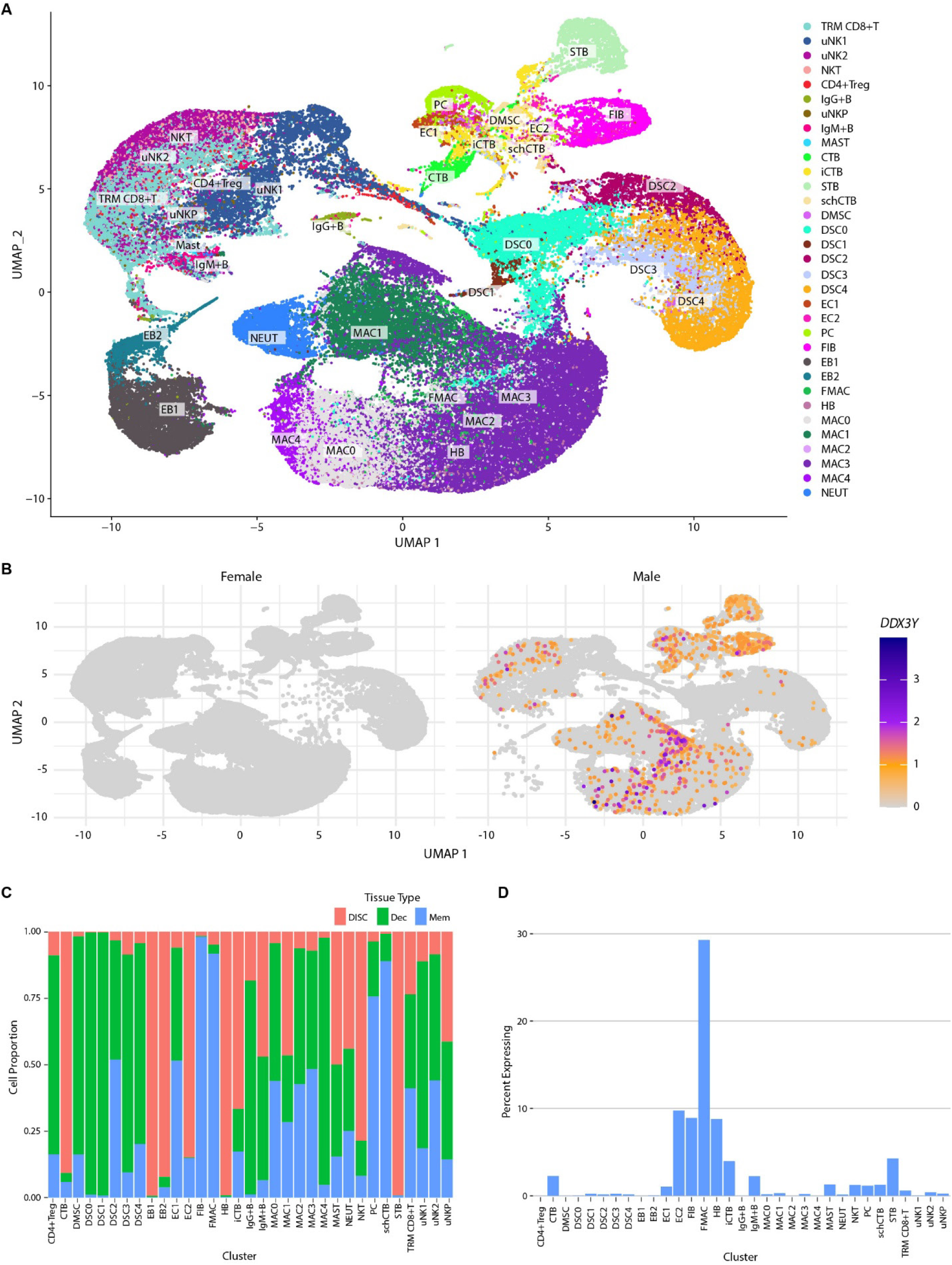
(A) The UMAP plot shows annotated cell populations in the late gestation pigtail macaque placenta. (B) A UMAP shows cells from placentas with ether a female (left) or male (right) fetus that expressed *DDX3Y*, a Y-chromosome gene. A yellow dot indicated that only a single cell in the cluster expressed *DDX3Y*, while a purple dot indicated that 3 or more cells expressed the gene. (C) Bar plots are shown with the percentage of cells from a cluster that originate from a specific tissue: the placental disc (DISC, red), chorioamniotic membranes (CAM, blue), and the maternal decidua (DEC, green). (D) Bar plots show the percent of *DDX3Y* expression in each cluster. Abbreviations: CTB, cytotrophoblast; DMSC, decidual mesenchymal stem cell; DSC, decidual stromal cell; EC, endothelial cells; EB, erythroblasts; FMAC, fetal macrophages; FIB, fibroblasts; HB, Hofbauer cells; iCTB, intermediate cytotrophoblast; IgM+B, IgM+ B cells; IgG+B, IgG+ B cells; MAC, macrophages; MAST, mast cells; NKT, natural killer T cells; NEUT, neutrophils; PC, pericyte; CD4+Treg, CD4+ regulatory T cells; schCTB, smooth chorion cytotrophoblast; STB, syncytiotrophoblast; TRM CD8+ T, tissue resident memory CD8+ Tcells; uNK, uterine natural killer cells; and uNKP, uNK precursor.

Annotation of cell populations considered gene expression profiles, as well as the unique attributes of each population related to their fetal origin and concentration within different placental tissues. We identified populations of uterine natural killer cells (uNK, 2 clusters), a uNK precursor (uNKP), tissue resident memory CD8+T cells (TRM CD8+ T cell, 1 cluster), regulatory CD4+ T cells (CD4+ Treg, 1 cluster), natural killer T cells (NKT,1 cluster), IgM+ and IgG+ B cells (2 clusters), maternal (MAC, 5 clusters) and fetal macrophages (FMAC, HB), neutrophils (NEUT, 1 cluster), erythroblasts (EB, 2 clusters), decidual mesenchymal stem cells (DMSC) and decidual stromal cells (DSC, 5 clusters), fibroblasts (FIB, 1 cluster), trophoblasts (3 clusters), endothelial cells (EC, 2 clusters), and pericytes (PC, 1 cluster).

Next, the origin of each population was determined by analyzing the expression profiles of *DDX3Y* in placentas with a male fetus. *DDX3Y*, also called *DBY*, encodes the DEAD-box RNA helicase and is located on the Y chromosome (Fig. 3B).(58) Next, the tissue origin of the cell populations was determined by leveraging our pipeline that processed and sequenced RNA transcripts separately from the DISC, CAM, and DEC (Fig. 3C). Similar to human placentas, the cell populations with a fetal origin included cytotrophoblast cells (CTB), intermediate CTB, and syncytiotrophoblast cells (STB) from the DISC, smooth chorion trophoblast cells from the CAM (schCTB), macrophages from the DISC (Hofbauer cells, HB), fibroblasts from the CAM (FIB), endothelial cells from the DISC (EC2), and a small macrophage population in the CAM (fetal macrophage, FMAC; Fig. 3D). Populations originating almost exclusively from the decidua included CD4+ regulatory T cells (CD4+ Treg), decidual mesenchymal stem cells (DMSC), decidual stromal cells (DSC0 and DSC1), and IgG+ B cells. *DDX3Y* expression was also noted in some immune cell and macrophage populations with a mixed tissue origin; these may be fetal cells originating from cord blood contained within the DISC. Due to small numbers, these were not sub-clustered and annotated. Although expression of *DDX3Y* was not observed frequently in the erythroblast populations (EB1, EB2), there were comparatively fewer of these cells isolated from male versus female placentas. Many cell populations had a mixed origin from DISC, CAM, and DEC, which may reflect the adherence of these tissues to one another. A population analogous to human extravillous trophoblast cells was not identified. Finally, a trajectory and latent time analysis was performed to reconstruct paths of cellular differentiation and to infer earlier and later stages of cellular differentiation for all cell populations in the placenta (Fig. 4A). This analysis predicted that the least differentiated populations were the DSC0, FIB, and uNKP populations, while the most differentiated were IgG+ B cells, EC1, STB, NEUT, and DSC3 (Fig. 4B, 4C). Overall, the identified cell populations in the pigtail macaque placenta were highly similar to those found in the human except for an absent extravillous trophoblast population, which was expected based on the literature.

**Figure 4.**
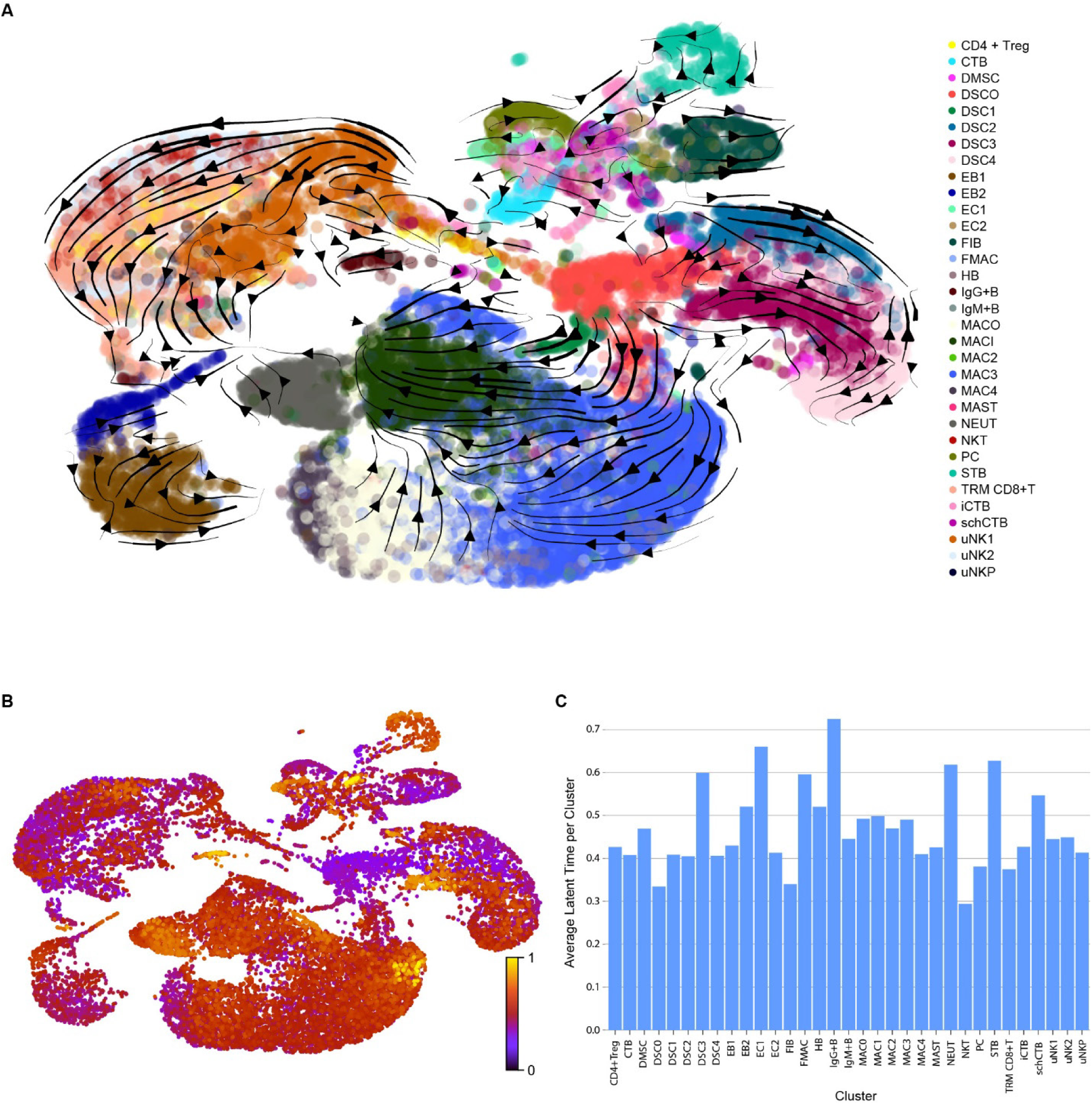
Trajectory and Latent Time Analyses. (A) The pseudotime of individual cells was inferred using trajectory analysis to predict the lineage, trajectory, and path of cells in the single-cell RNA-Seq atlas of the pigtail macaque placenta. (B) A latent time analysis was performed to estimate the real time experienced by cells as they differentiate based on the cells’ transcriptional dynamics. Purple and yellow colors indicate a less and more differentiated population, respectively. (C) A bar plot shows the average latent time elapsed for each cluster. Higher numbers indicate an older inferred age of the cell population.

### Trophoblast, Pericyte, Endothelial Cell, and Fibroblast Clusters

We identified three populations of trophoblast cells originating from the DISC and one from CAM (UMAP, Fig. 5A). Cytotrophoblasts (CTB), intermediate CTB (iCTB), syncytiotrophoblasts (STB), and smooth chorion cytotrophoblasts (schCTB) were identified in the pigtail macaque placenta based on expression of canonical genes like GATA binding protein 3 (*GATA3*), transcription factor AP-2 gamma (*TFAP2C*), and keratin 7 (*KRT7*). A subcluster annotated as CTB expressed canonical genes associated with human CTB including chorionic somatomammotropin hormones (*CSH2*, *CSH3*, *CSH4*), corticotropin releasing hormone (*CRH*), serpin family E member 2 (*SERPINE2*), placenta-specific 8 (*PLAC8*), vascular endothelial growth factor receptor 1 (*VEGFR1* or *FLT1*), *CD36*, and the integrin Subunit Alpha 5 (*ITGA5*; Fig. 5B). The CTB populations (schCTB, CTB, iCTB) expressed canonical markers including paternally expressed 3 (PEG3) and serine peptidase inhibitor, Kunitz type 1 (*SPINT1*) that differentiated them from the STB population. They also highly expressed endogenous retrovirus group 48 member 1, envelope (*ERVH48-1*, ENSMMUG00000044879), which encodes suppressyn, a protein that inhibits syncytialization of villous cytotrophoblasts.(59) We did not anticipate finding a large population of STB, as the 10X Genomics Chromium platform does not efficiently capture large multinucleated STB populations. However, we identified a population of cells that expressed canonical STB markers. This population expressed the endogenous retrovirus group FRD member 1 (*ERVFRD-1*), which encodes the syncytin-2 protein crucial for the fusion of trophoblast cells into the syncytium, and syndecan 1 (*SDC1*).(60) The STB population also expressed key STB differentiation markers including insulin-like 4 (*INSL4*), PDZ and LIM domain 2 (*PDLIM2*), and CCAAT enhancer binding protein beta (*CEBPB*). Several genes involved in hormone secretion and are characteristic of STBs were also upregulated in this population: cytochrome P450 family 19 subfamily A member 1 (*CYP19A1*), chorionic gonadotrophin subunit alpha (*CGA*), corticotropin releasing hormone (*CRH*), chorionic somatomammotropin hormone 2 (*CSH2*), chorionic somatomammotropin hormone 3 (*CSH3*), and chorionic somatomammotropin hormone 4 (*CSH4*). Finally, a trophoblast population originating from the CAM was called schCTB and expressed canonical markers of the corresponding human population including: *LAMB3*, *LAMC2*, and *KRT5* (Fig. 5B).

**Figure 5.**
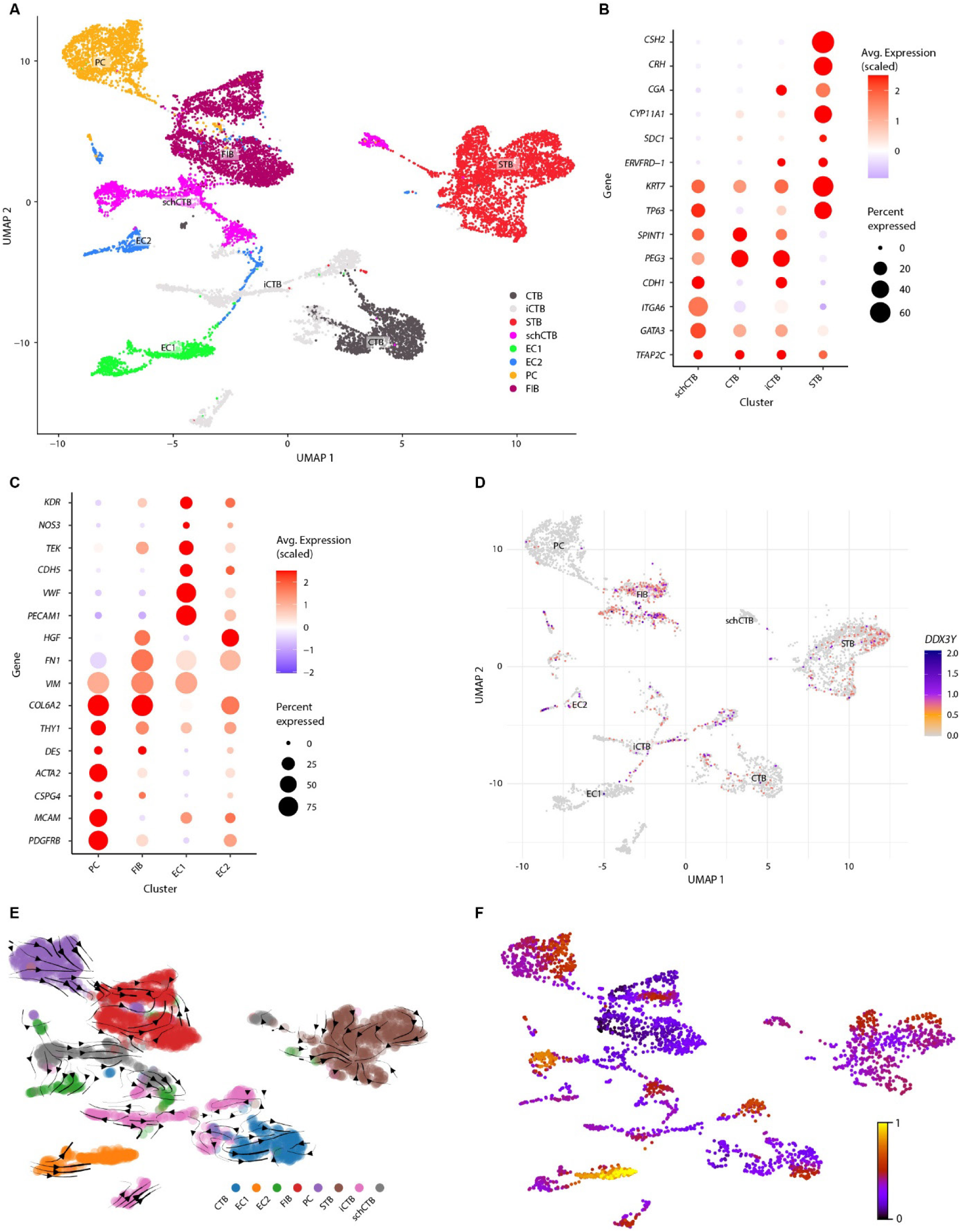
Trophoblast, pericyte, endothelial, and fibroblast cells. (A) Trophoblast, PC, EC, and FIB populations were separately subclustered from other populations and shown in a UMAP. (B, C) Dot plots show canonical gene markers. The scaling of colors and dot size is with respect to all genes from all cell populations. (D) A UMAP of cells is shown reflecting *DDX3Y* gene expression in male placentas. (E) The trajectory analysis shows RNA velocity with black lines representing the inferred direction and speed of cell state transitions. Arrows on the plot indicate the predicted future state of cells, with their direction pointing towards the estimated trajectory. Arrow length corresponds to the magnitude of velocity, reflecting the rate of change in gene expression. (F) Cells were colored by latent time, from early (dark purple) to late (yellow) stages of differentiation. Abbreviations: CTB, cytotrophoblast; EC, endothelial cells; FIB, fibroblast; iCTB, intermediate cytotrophoblast; PC, pericyte; schCTB, smooth chorion cytotrophoblast; and STB, syncytiotrophoblast.

On the UMAP, the PC, EC, and FIB were in close proximity to one another (Fig. 5A). Dot plots of gene expression in each population were consistent with gene profiles of the same populations in humans (Fig. 5B, 5C). As expected in a largely differentiated third-trimester placenta, the trajectory analysis of RNA velocity revealed only a limited connection between populations (Fig. 5C). Arrows connected the iCTB and CTB subclusters; directionality of the arrows indicated that iCTB was a possible progenitor cell for the CTB cluster but might also represent an intermediate stage of CTB differentiation. Neither the iCTB nor the CTB subclusters were directly connected with the STB subcluster. The iCTB subcluster expressed *AGAP1* and *ADORA2B*, key genes involved in trophoblast development (Fig. 5B). Nearly all subclusters contained fetal cells except EC1; rare cells in the PC population were of fetal origin (Fig. 5D). The trajectory analysis indicated movement from the PC into the FIB subclusters, two stromal cell populations originating predominantly from the CAM (Fig. 5E). The trajectory of pseudotime in the iCTB population moved into the CTB cluster. A latent time analysis considering only these populations indicated that the maternal EC1 population was the most differentiated (Fig. 5F).

### uNK, T Cell and B Cell Clusters

After initial unsupervised clustering, the T and uNK cell populations were difficult to annotate; therefore, the large immune cell cluster was reclustered separately at a higher resolution (0.25; Fig. 6A). After this step, distinct T cell and uNK cell populations were identified based on canonical markers defined in the literature (Fig. 6B). A large population of T cells emerged with the largest expressing *CD8A* and *IL7R*, which is analogous to the expression of TRM CD8+ T cells in human placentas (Fig. 6B). A smaller T cell population expressed *CD4*, *FOXP3* and *CD25*, canonical markers for a CD4+ Treg population (Fig. 6B). NKT cells were also identified by their co-expression of *CD8A* (CD8; Fig. 6B) and *FCGR3* (CD16, Fig. 6C). Two uterine NK (uNK) cell populations and a precursor were defined by their gene expression [*NCAM1+* (CD56+), *ITGAM+*, *CD3D-*, *FCGR3-*(CD16-)], similar to populations previously described (38) uNK1 cells were characterized by their cytotoxic granule gene expression, such as PRF1, *GNLY*, *GZMA*, and *GZMB*; uNK2 cells expressed *GZMK*, *GZMM*, and *XCL1* (Fig. 6C). Two B cell populations were identified expressing IgM and IgG heavy chains, and a small mast cell cluster (Fig. 6D). Sparse fetal cells were detected with a small cluster co-located within the CD4+ Treg population (Fig. 6E). A trajectory analysis indicated movement between the uNKP and uNK1 or uNK2 populations (Fig. 6F). It was unclear whether this directionality indicated that the uNKP was a true precursor of both populations, or an intermediate stage between the two populations. Both the uNK2 and CD8+ TRM populations appeared to be a source for NKT cells (Fig. 6F). The latent time analysis indicated that the CD4+ Treg population was the most differentiated of these populations (Fig. 6G).

**Figure 6.**
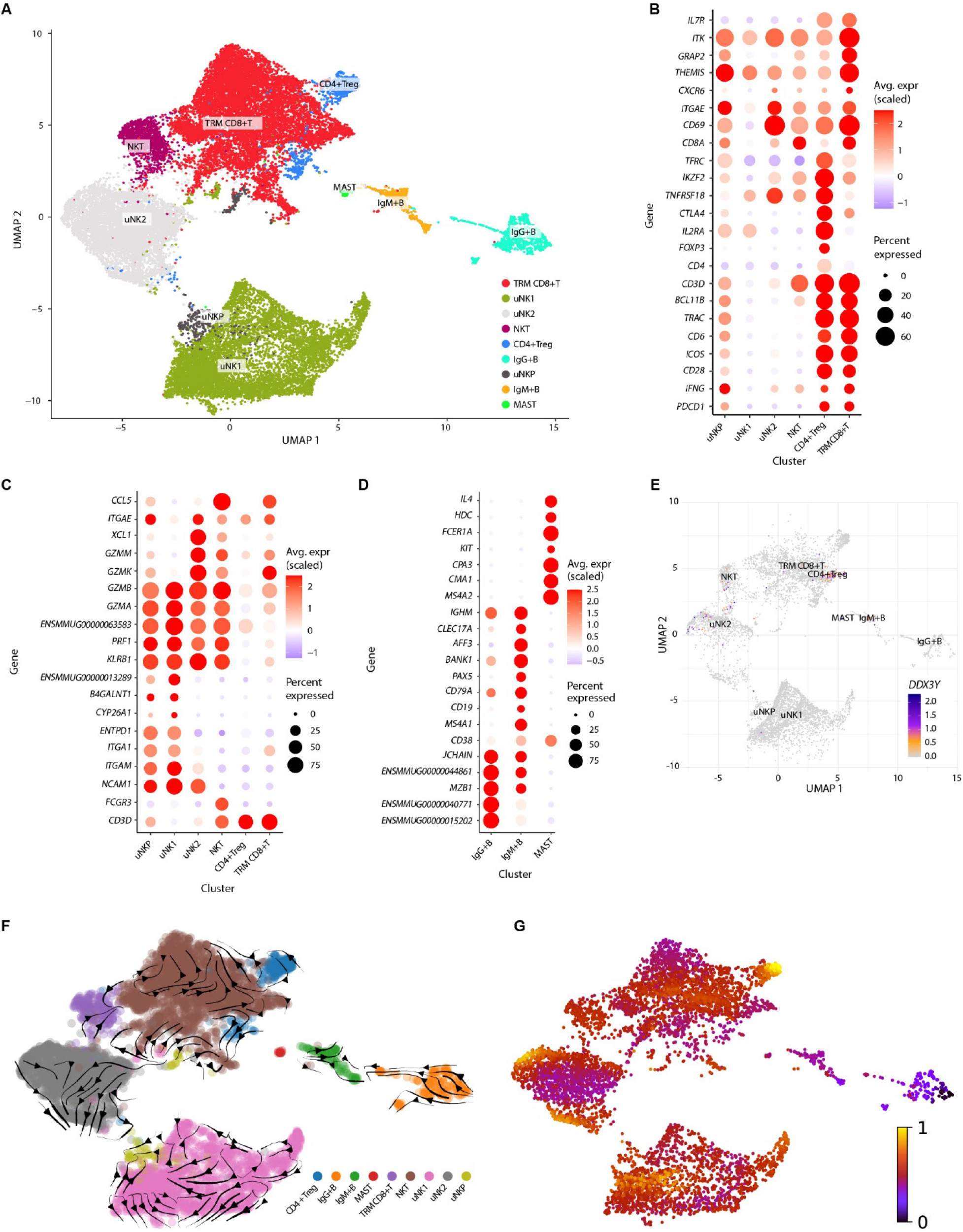
uNK, T cell and B cells. The uNK, T cell, and B cell populations were visualized in a UMAP (A) and represented in a dot plot (B, C, D) using canonical gene markers. The average expression of genes shown in the plot is scaled with respect to all genes from all cell populations. (E) A UMAP of cells is shown reflecting *DDX3Y* gene expression in male placentas. (F) A trajectory analysis shows RNA velocity across the uNK, T, and B cell populations with black lines representing the inferred direction and speed of cell state transitions. Arrows on the plot indicate the predicted future state of cells, with their direction pointing towards the estimated trajectory. Arrow length corresponds to the magnitude of velocity, reflecting the rate of change in gene expression. (G) Cells were colored by latent time, from early (dark purple) to late (yellow) stages of differentiation. Abbreviations: CD4+Treg, CD4+ regulatory T cells; ENSMMUG00000013289, homolog is the leukocyte immunoglobulin-like receptor, subfamily B, member 1 gene (*LILRB1*); ENSMMUG00000015202, homologs include immunoglobulin heavy chain genes (IGHG1/IGHG2/IGHG3/IGHG4); ENSMMUG00000040771, homolog is the immunoglobulin heavy constant gamma 2 gene (*IGHG2*); ENSMMUG00000044861, homologs include antibody light chain genes (*IGLC1*/*IGLC2*/*IGLC3*/*IGLC7*/*IGLL5*); ENSMMUG00000063583, homolog is granulysin (*GNLY*); IgG+B, IgG+ B cells; IgM+B, IgM+ B cells; MAST, mast cells; NKT, natural killer T cells; TRM CD8+ T, tissue resident memory CD8+ Tcells; uNK, uterine natural killer cells; and uNKP, uNK precursor.

### Macrophages and Neutrophils

Diverse macrophage sub-populations within the pigtail macaque placenta were captured by the identification of two fetal macrophage clusters (HB, FMAC), and five maternal macrophage clusters (MAC0, MAC1, MAC2, MAC3, MAC4; Fig. 7A). The MAC1 population exhibited a pro-inflammatory M1 phenotype and expressed *CD14*, *FCGR3* (CD16), interleukin-1 beta (*IL1B*), and thrombospondin-1 (*THBS1*; Fig. 7B). In contrast, the MAC0 population exhibited an expression profile more characteristic of anti-inflammatory M2 macrophages with poor expression of CD14 and CD16 (Fig. 7B). MAC2 highly expressed C-C Motif Chemokine Ligand 17 (*CCL17*) and the gene that encodes part of the major histocompatibility complex (MHC) class II system in rhesus macaques (*MAMU-DRB1*; Fig. 7B). Gene expression of MAC3 was intermediate between MAC0 and MAC1. A population of neutrophils (NEUT) was also identified, which expressed the protein tyrosine phosphatase receptor Type C (*PTPRC* encoding CD45), *CD14*, colony-stimulating factor 3 receptor (*CSF3R*), L-selectin (*SELL* encoding CD62L), and carcinoembryonic antigen-related cell adhesion molecule 8 (CEACAM8 encoding CD66b; Fig. 7B). Identification of the HB and FMAC populations from the DISC and the CAM, respectively, leveraged the *DDX3Y* analysis in male placentas (Fig. 7C). The trajectory analysis identified movement from the MAC3 population into both MAC0 and MAC1, and from MAC4 into MAC0 (Fig. 7D). A latent time analysis inferred that MAC2 was the most differentiated population of these clusters (Fig. 7E).

**Figure 7.**
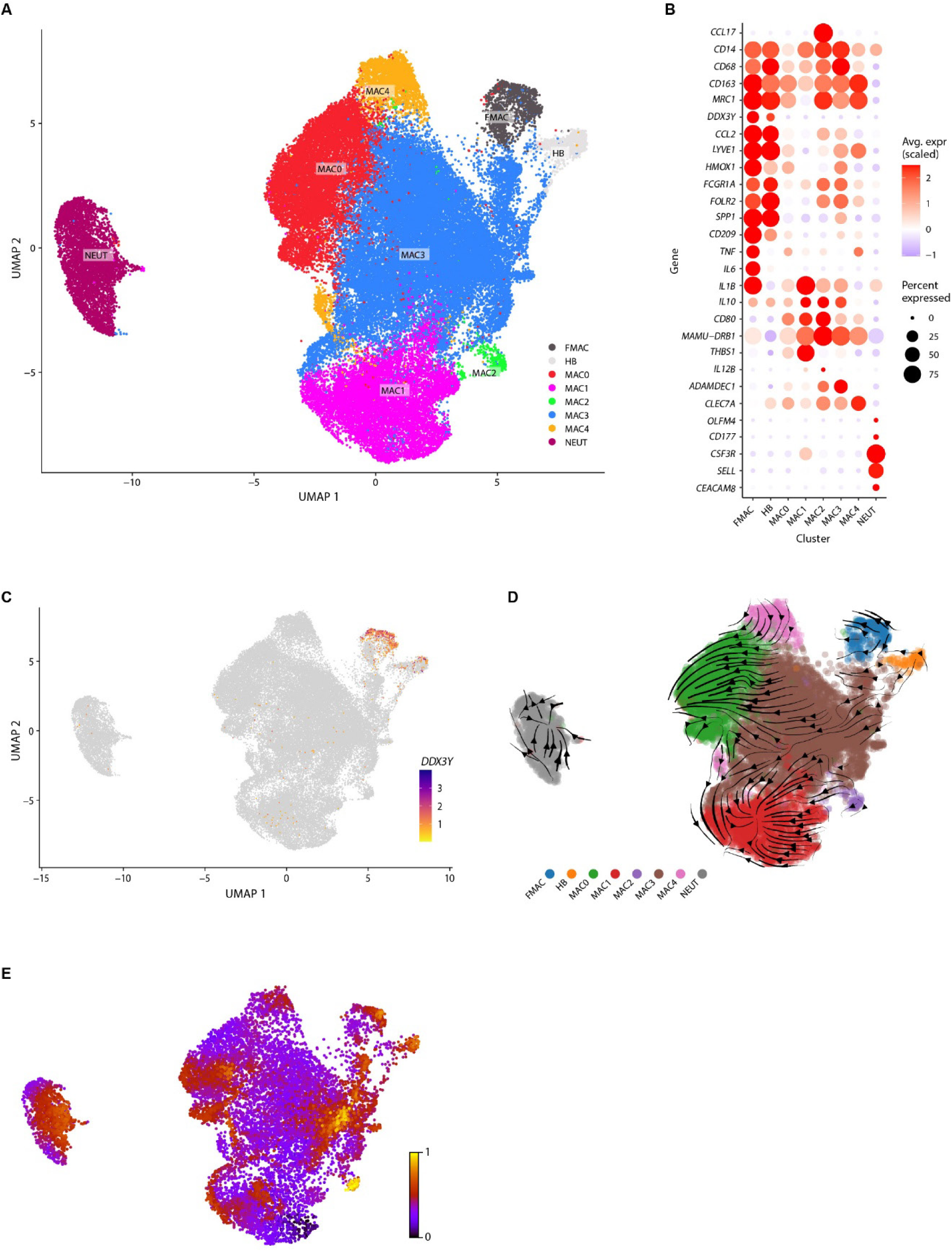
Macrophage populations. (A) The macrophage cell populations were visualized in a UMAP. (B) A dot plot is shown to depict canonical and distinguishing gene markers across populations. The average expression of genes shown in the plot is scaled with respect to all genes from all cell populations. (C) A UMAP of cells is shown reflecting *DDX3Y* gene expression in male placentas. (D) A trajectory analysis shows RNA velocity with lines representing the inferred direction and speed of cell state transitions. Arrows on the plot indicate the predicted future state of cells, with their direction pointing towards the estimated trajectory. (E) Cells were colored by latent time, from early (dark purple) to late (yellow) stages of differentiation. Abbreviations: DMSC, decidual mesenchymal stem cell; FMAC, fetal macrophage; HB, Hofbauer cells; MAC, macrophage; and NEUT, neutrophil.

### Fetal Erythroblast Cells

The placenta is a hematopoietic organ, where primitive fetal red blood cells mature and enucleate.(61) We identified two fetal erythroblast (EB) populations, EB1 and EB2, which were located primarily in the DISC (Fig. 8). EB1 and EB2 were characterized by high expression of hemoglobin subunit genes, including hemoglobin subunit alpha (*HBA*), hemoglobin subunit epsilon 1 (*HBE1*), and hemoglobin subunit Mu (*HBM*). Both EB1 and EB2 also expressed *CD71*, a potent immunoregulatory gene. EB2 represented a more mature population with expression of glycophorin A (*GYPA*) and phosphoethanolamine/phosphocholine phosphatase 1 (*PHOSPHO1*).

**Figure 8.**
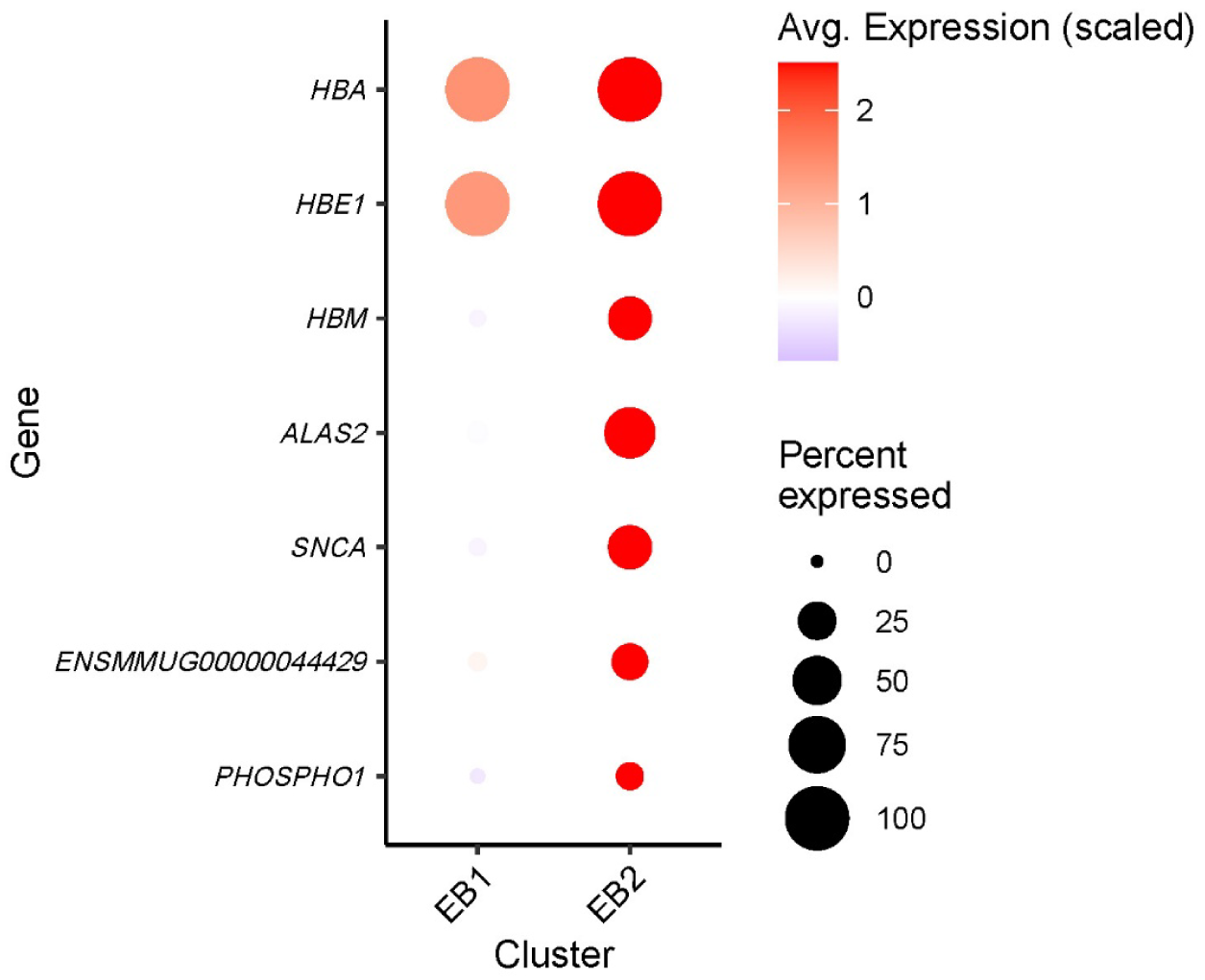
Fetal Erythroblast Cells. Canonical gene markers for the two fetal EB populations are shown. Abbreviations: EB, erythroblast; ENSMMUG00000044429, GYPA.

### Decidual Stromal Cells

Decidual stromal cells play an essential role in the facilitation of successful implantation and maintenance of pregnancy. Six decidual stromal cell (DSC) populations were identified by expression of canonical markers such as periostin (*POSTN*), tissue inhibitor of metalloproteinases 3 (*TIMP3*), and insulin-like growth factor-binding protein 1 (*IGFBP1*; Fig. 9A). The decidual mesenchymal stem cell (DMSC) population expressed key stem cell markers, including Thy-1 cell surface antigen (*THY1*), Vimentin (*VIM*), 5’-nucleotidase ecto (*NT5E*; Fig. 9B). A gradient of gene expression suggests the DSC populations were captured at various stages of progesterone-induced decidualization, transitioning from the least decidualized (DMSC) to more decidualized populations (DSC2, DSC3, and DSC4; Fig. 9B). The profile of RNA expression for DSCO and DSC1 was more consistent with less decidualized DSCs, based on their lack of expression of prolactin (*PRL*) and the progesterone receptor (*PGR*; Fig. 9B). As expected, fetal cells were rare in this population (Fig. 9C). The trajectory analysis inferred movement of multiple populations into DSC4, including DMSC, DSC3 and DSC2 (Fig. 9D). Interestingly, the origin of DSC0 and DSC1 appeared to be DSC3 and DSC4, suggesting dedifferentiation of these populations into a less decidualized state (Fig. 9D). The latent time analysis predicted that DSC3 was the most differentiated of these populations (Fig. 9E).

**Figure 9.**
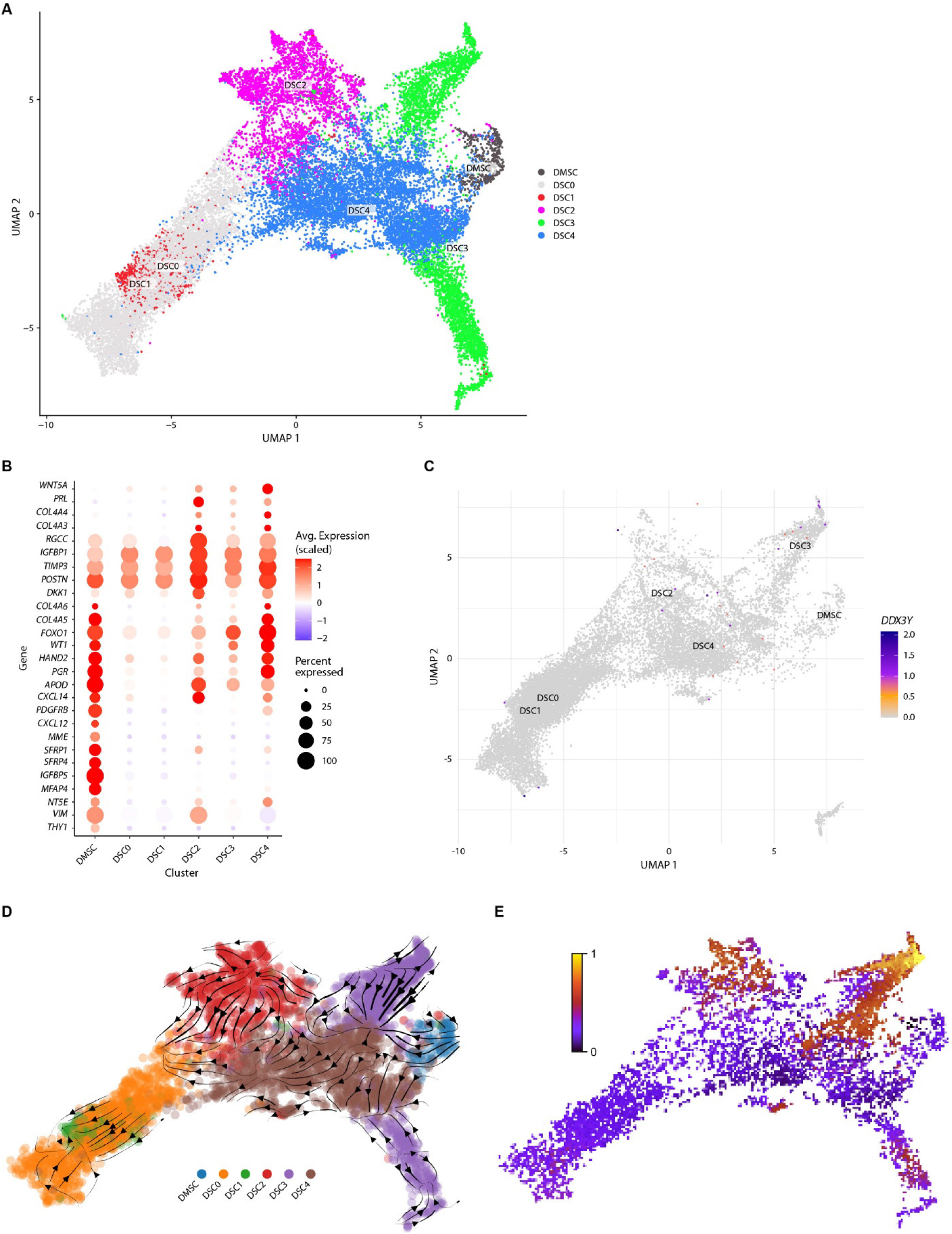
Decidual stromal cell (DSC) populations. (A) The DSC populations were visualized in a UMAP. (B) The dot plot shows gene markers defining decidual mesenchymal stem cells (DMSC) and DSC. The average expression of genes shown in the plot is scaled with respect to all genes from all cell populations. (C) A UMAP of cells is shown reflecting *DDX3Y* gene expression in these clusters from pregnancies with male placentas. (D) A trajectory analysis shows RNA velocity with lines representing the inferred direction and speed of cell state transitions. Arrows on the plot indicate the predicted future state of cells, with their direction pointing towards the estimated trajectory. (E) Cells were colored by latent time, from early (dark purple) to late (yellow) stages of differentiation.

## Discussion

We describe a scRNA-Seq atlas of the pigtail macaque placenta for the first time, which was deeply sequenced using tissues isolated separately and then aggregated from the placental basal plate, chorioamniotic membranes, and maternal decidua. This approach allowed more specific annotation of stromal trophoblast and macrophage populations reflecting their tissues of origin. We found a diversity of uterine NK cells, stromal cells, and macrophages that are nearly identical to those described at the human maternal-fetal interface. A notable difference with the human placenta was the absence of extravillous trophoblast cells, which has been previously reported in several macaque species.(31) Overall, the similarities in placental architecture and cellular composition suggest that the pigtail macaque is a robust model of the human placenta.

NHP models are a valuable tool for determining disease pathogenesis of reproductive disorders and for trialing novel therapeutics. A scRNA-Seq atlas of the normal pigtail macaque placenta provides a detailed reference against which we can evaluate how experimental manipulations change the cellular composition and gene expression at the maternal-fetal interface. Scientists studying species of closely related nonhuman primates, like the rhesus macaque (Macaca mulatta) and crab-eating macaque (Macaca fascicularis), can also use this atlas to accelerate identification of single-cell populations from the placentas of their own models.

Uterine natural killer (uNK) cells, macrophages, and extravillous trophoblast cells are collectively responsible for transforming uterine spiral arteries into wide vessels with lower resistance to blood flow. The two uNK cell clusters in the pigtail macaque placenta were similar to both the human uNK1 (high expression of cytotoxic granules) and uNK2 (*XCL1* expression). A third uNK population described in the human first-trimester placenta, uNK3, was not identified in the late gestation pigtail macaque placenta. The human uNK3 population expresses CCL5 and is thought to aid in regulating extravillous trophoblast invasion. Interestingly, in human placentas, the uNK1 and uNK2 cells predominate in the first trimester, while uNK3 cells are most common in the third trimester.(62) The absence of the uNK3 population in the pigtail macaque may also contribute to the shallower trophoblast invasion of the spiral arteries.(2)

The strength of this resource lies in the deep sequencing of many tissues at the maternal-fetal interface and the careful annotation of diverse cell types, which has yielded one of the most complete atlases of the maternal-fetal interface in any species. One example of the richness of this atlas is in the identification of two erythroblast populations, which reflects the capture of nucleated, immature red blood cells in the fetal blood vessels present in the placental tissues. Another strength is in the trajectory and latent time analyses, which allowed evaluation of the relationships of populations to one another and their relative differentiation. This resource will also enable the deconvolution of bulk RNA-Seq datasets to estimate the proportion of single cells without the investment required in building a single-cell RNA-Seq dataset. Limitations of the work include an incomplete annotation of the pigtail macaque genome, which necessitated using the closely related rhesus macaque genome as a reference. Even using the rhesus macaque genome, there were genes expressed in the placenta that we could not identify; however, this limitation did not prevent cell identification, which was evident from the RNA profile as a whole. An ongoing effort is needed to improve the annotation of these genomes and reflect the genetic diversity across these species.

In summary, this single-cell atlas of the late pigtail macaque placenta provides a valuable resource for understanding both normal pregnancies and experimental NHP models of reproductive disorders. The pigtail macaque is a scientifically important species for understanding normal pregnancy and adverse pregnancy outcomes, including preterm labor and the maternal-fetal response to infectious disease.(5, 14, 23, 26, 27, 50–52, 63, 64) As the pigtail macaque is evolutionarily very close to the rhesus and cynomolgus macaques, this atlas may have translational value for multiple species in which there are scientific models of reproductive disorders.

## Materials and Methods

### Ethics Statement

All nonhuman primate experiments were performed in strict accordance with the recommendations in the Guide for the Care and Use of Laboratory Animals of the National Research Council and the Weatherall report, “The use of non-human primates in research.” The Institutional Animal Care and Use Committee (IACUC) of the University of Washington (UW) approved the study’s protocols (4165-01, last approval date: 05/07/2025; 4165-02, last approval date: 04/09/2025). All surgeries were performed under general anesthesia, and all efforts were made to minimize pain and distress.

### Study design

Placentas and decidua were obtained from ten pigtail macaques enrolled as uninfected controls on two different experimental protocols and thus, had two different experimental manipulations (Table S1). One control group received choriodecidual and intra-amniotic saline infusions after surgical implantation of choriodecidual and amniotic catheters, which were compared to experimental infections with Group B Streptococcus (n=5). A second group received five subcutaneous inoculations of media along the forearm to serve as controls for a study of Zika virus pathogenesis (n=5). The gestational age at tissue collection spanned the late second and third trimester (118-150 days gestation; term gestation = 172 days). After cesarean section, maternal decidua and placental tissues (basal plate and chorioamniotic membranes) were separated and collected for single cell dissociation and RNA-Seq analysis.

### Single Cell RNA-Seq and Bioinformatics Pipeline Single Cell Dissociation

#### Isolation of cells from chorionic villous tissue of the placental disc (DISC) and maternal decidual tissues (DEC)

Cell dissociation from placental tissue was performed following a previously described protocol.[39] Decidual and chorionic villous tissues were finely chopped into approximately 0.2 mm³ cubes using sterile scissors. The tissue fragments were enzymatically digested in 15 mL of a 0.4 mg/mL collagenase V solution (Sigma, C-9263) prepared in working medium, in 50 mL conical vials. The working medium consisted of RPMI 1640 (Thermo Fisher Scientific, 21875-034), supplemented with 10% heat-inactivated fetal bovine serum (FBS; Thermo Fisher, SH3007103), 50 U/mL penicillin-streptomycin (Pen-Strep; Thermo Fisher, 15-140-122), and 250 U/mL DNase I (Sigma, D4527-20KU). The digestion was carried out at 37 °C for 45-60 minutes with continuous stirring using a magnetic stir bar on a heated plate.

Following digestion, the supernatant was sequentially filtered through 100-μm and 40-μm cell strainers (Fisher Scientific, 22-363-549 and 22-363-547, respectively). The filtered cell suspension was centrifuged at 300 × g for 5 minutes at room temperature. The resulting pellet was resuspended in 5 mL of red blood cell lysis buffer (RBC) (Invitrogen, 00-4300-54) and incubated for 10 minutes at room temperature to lyse residual erythrocytes. For wash steps, 1x phosphate-buffered saline (PBS) containing 2% FBS was added to samples that contain RBC lysis buffers and washed twice. After RBC lysis, the samples were washed twice with 1× (PBS) containing 2% fetal bovine serum (FBS) to remove residual lysis buffer and cellular debris. Cells were then counted, and the cell viability was assessed using trypan blue.

#### Isolation of cells from the chorioamniotic membranes (CAM)

Each CAM was placed in a sterile Petri dish, and clogs or debris were gently removed using scissors under 1× phosphate-buffered saline (PBS). The cleaned membrane tissue was enzymatically digested in 70 mL of a digestion solution consisting of 0.2% Trypsin 250 (Pan Biotech, P10-025100P), 0.02% EDTA (Sigma, E9884), and 5 U/mL Dispase (Corning, 354235) in 1× Dulbecco’s Modified Eagle Medium (DMEM), with stirring at 37 °C for 10 minutes with continuous stirring using a magnetic stir bar on a heated plate. The disaggregated cell suspension was passed through sterile food-grade muslin gauze (Winware). The tissue was washed through the gauze using working medium composed of RPMI 1640 (Thermo Fisher Scientific, 21875-034), supplemented with 10% heat-inactivated fetal bovine serum (FBS; Thermo Fisher, SH3007103), 50 U/mL penicillin-streptomycin (Pen-Strep; Thermo Fisher, 15-140-122), and 250 U/mL DNase I (Sigma, D4527-20KU). The resulting filtrate was collected and set aside. The undigested gelatinous tissue retained on the gauze was recovered and subjected to a second enzymatic digestion using 25 ml of 0.4 mg/mL collagenase V (Sigma, C9263) in the working medium. This digestion was performed with gentle shaking at 37 °C for 30 minutes. The resulting cell suspension was again passed through sterile muslin gauze, and cells were pelleted from the filtration by centrifugation as previously described.

Cells obtained from both the trypsin/dispase and collagenase digestions were pooled and sequentially filtered through 100-μm and 70-μm cell strainers (Fisher Scientific, 22-363-549 and 22-363-547, respectively), followed by a wash in Ham’s F-12 medium. The filtered cell suspension was centrifuged, and the pellet was resuspended in 5 mL of red blood cell lysis buffer (Invitrogen, 00-4300-54) and incubated at room temperature for 10 minutes. After RBC lysis, the samples were washed twice with 1× phosphate-buffered saline (PBS) containing 2% fetal bovine serum (FBS) to remove residual lysis buffer and cellular debris. Cells were then counted, and cell viability was assessed using trypan blue.

### Sequencing

10x Chromium single-cell libraries were prepared according to the standard protocol outlined in the manual (Chromium Next GEM Single Cell 3’ kits v3.1: dual index, 10X Genomics, Pleasanton, CA). Briefly, we loaded single-cell suspensions, 10x barcoded gel beads, and reagents onto a “Chromium Single Cell A Chip” to generate Gel Bead-In-EMulsions (GEMs) using the 10X Chromium Controller (10X Genomics, Pleasanton, CA). Reverse transcription of mRNA occurred inside each GEM, followed by amplification of cDNA and preparation of sequencing libraries according to manufacturer instructions. Quality control was performed in multiple steps to check the quality of the cDNA and the library using the Agilent 2200 TapeStation system (Agilent Technologies, Santa Clara, CA). Finally, shallow sequencing was performed using a Next-Seq 2000 (Illumina Inc., San Diego, CA) to evaluate the percentage of reads mapping to the reference genome, GC content distribution, and sequencing depth (number of reads per cell). Based on this information, we determined if the cDNA library was of sufficient quality to send for deep sequencing, and whether an adjustment to the amount of any samples was necessary to achieve a greater number of reads per cell. Deep sequencing of the cDNA library was performed on a Nova-Seq (Illumina Inc., San Diego, CA) to achieve a read depth of approximately 40,000 reads per cell.

### Genome Alignment

Raw FASTQ files were processed using Cell Ranger v7.2.0 software (10x Genomics, Pleasanton, CA, USA). Reads were initially aligned to the pigtail macaque (*M. nemestrina)* genome (Mnem_1.0, GenBank Assembly ID: GCA_000956065.1, 2015). Initial analysis revealed that key canonical genes of interest to define placental cell types were not annotated in the *M. nemestrina* genome, which is a lower quality (scaffold-based) genome. In contrast, the rhesus macaque genome (Mmul_10, GenBank Assembly ID: GCA_003339765.3) has a contiguous assembly and is more completely annotated than the pigtail macaque. As the rhesus and pigtail macaque are closely related species, alignment was performed to the rhesus macaque genome.

### Quality Control, Normalization, and Harmonization

Next, samples were aggregated with Cell Ranger. Empty droplets and potential doublets were removed by filtering out cells with fewer than 200 genes or more than 2500 genes. Damaged cells were removed by filtering out cells with greater than 3% mitochondrial reads. With the Seurat package in R (Seurat version 5.1.0, R version 4.4.1)(65–69), the data set was normalized with the SCTransform(70) function and then the first 100 principal components were found with the RunPCA function. Data was corrected for batch effects using the Harmony package (version 1.2.0)(71) and the top 30 harmony components were processed using the Seurat runUMAP function to embed and visualize the cells in a two-dimensional map via the Uniform Manifold Approximation and Projection for Dimension Reduction (UMAP) algorithm. A resolution of 0.25 was used to cluster the single cells, resulting in 33 clusters. Next, scDEED was used to detect dubious cell embeddings within the default UMAP settings and then to optimize UMAP parameters to minimize distortion.(72) scCustomize was then used to generate final UMAPs.(73)

### Pseudobulking

Data from the per-sample count matrices were imported into R and analyzed with edgeR (v3.38.1) under Bioconductor (v3.16). We compared the uninfected media and saline control groups using pseudobulking by exploring the data by calculating the biological coefficient of variation (BCV). The BCV was defined as the square root of the tagwise dispersion. A generalized linear model was fitted by contrasting the saline and medical controls was performed using the R function, glmLRT(coef = 2).

### Cell Clustering, Annotation, and Identification of Fetal Cells

After investigation of the top genes expressed by each cluster, it was determined that some clusters required further refinement. Reclustering was performed separately on the B/T/NK and monocyte/macrophage clusters using a resolution of 0.25 in Seurat. Canonical markers were then used to annotate new clusters and the new cell population labels were integrated back into the main UMAP. To distinguish between fetal and maternal cells within the atlas, we evaluated the distribution of *DDX3Y* expression, a Y-chromosome gene, in placentas from pregnancies with a male fetus.(58)

### Trajectory Inference

We used the Python 3(74) packages scVelo(75) and Velocyto(76) to reconstruct the trophoblast, stromal cell, and macrophage lineages from our single-cell gene expression data. This method utilized a likelihood-based dynamic model to characterize the RNA velocity of cell subpopulations and to infer the trajectory of cellular age.

## Acknowledgment

We are grateful to Riley Raker for assistance with graphic design.

## Data Availability

The data is available in the NCBI Gene Expression Omnibus (GSE305531). Bioinformatics code is available on the <u>Adams Waldorf Lab GitHub</u>.

## Author Contributions

All authors listed have made a substantial, direct, and intellectual contribution to the work and approved it for publication.

## Supporting Information Figure Legends

**Figure S1.** Proportion of Single-Cell Populations by Type of Control Group. This figure illustrates the proportion of single-cell populations (y-axis) out of all single-cell populations for each type of control: subcutaneous media inoculation (CTRL) or choriodecidual saline (SAL) inoculation. Each dot within a bar plot represents a single sample from either the placental basal plate, chorioamniotic membranes or maternal decidua. The black box shows the median and the interquartile range. The blue bounding box indicates the mean +/- one standard deviation. A red colored box indicates a significant difference in proportions between the groups using a Fisher’s exact test (p<0.05). Abbreviations for single-cell populations are shown in Figure 3.

**Figure S2.** Proportion of Single-Cell Populations by Tissue of Origin. This bar plot illustrates the proportion of single-cell populations (y-axis) among all single-cell populations from each tissue source: placental disc (DISC), maternal decidua (DEC), and chorioamniotic membranes (CAM). Each dot within the bar plot represents a single sample from either the media or saline control groups. The black box shows the median and the interquartile range. The blue bounding box indicates the mean +/- one standard deviation. A colored box indicates a significant difference in proportions between either: 1) the DEC (red) and the CAM, or 2), the DISC (turquoise) and the CAM, using a constrained beta-binomial distribution (p<0.05). Abbreviations for single-cell populations are shown in Figure 3.

**Figure S3.** Proportion of Single-Cell Populations by Fetal Sex. This bar plot illustrates the proportion of single-cell populations (y-axis) among all single-cell populations from each placenta group from a single sex: female (F) versus male (M). Each dot within the bar plot represents a single sample from either the media or saline control groups from a specific tissue (basal plate, membranes, decidua) grouped by fetal sex. The black box shows the median and the interquartile range. The blue bounding box indicates the mean +/- one standard deviation. A red colored box indicates a significant difference in proportions between placentas from a female versus a male fetus using a Fisher’s exact test (p<0.05). Abbreviations for single-cell populations are shown in Figure 3.

