## Supplemental Information for "A single-cell transcriptomic atlas of the pigtail macaque placenta in late gestation"

### Table of Contents

TABLE S1. ANIMAL DEMOGRAPHICS ---------------------------------------------------------------------- 2

FIGURE S1. PROPORTION OF SINGLE-CELL POPULATIONS BY TYPE OF CONTROL GROUP ---------------------------------------------------------------------------------------------------------- 3

FIGURE S2. PROPORTION OF SINGLE-CELL POPULATIONS BY TISSUE OF ORIGIN------ 4

FIGURE S3. PROPORTION OF SINGLE-CELL POPULATIONS BY FETAL SEX ---------------- 5

**Table S1.** Animal Demographics

| Animal ID | Inoculum | Maternal Age at Delivery (years) | Gestational Age at Inoculation (days) | Gestational Age at Delivery (days) | Fetal Body Weight (grams) | Fetal Sex |
|---|---|---|---|---|---|---|
| CTRL01 | Media | 5.23 | 138.00 | 158 | 499.2 | Female |
| CTRL02 | Media | 13.49 | 132.38 | 155 | 573.0 | Male |
| CTRL03 | Media | 12.37 | 133.38 | 154 | 411.9 | Female |
| CTRL04 | Media | 10.76 | 127.38 | 147 | 469.0 | Male |
| CTRL05 | Media | 12.88 | 141.34 | 144 | 412.6 | Male |
| CTRL06 | Saline | 11.26 | 134.36 | 138 | 337.6 | Female |
| CTRL07 | Saline | 5.68 | 127.32 | 128 | 242.0 | Male |
| CTRL08 | Saline | 6.05 | 133.38 | 134 | 264.0 | Female |
| CTRL09 | Saline | 14.74 | 135.31 | 136 | 299.5 | Female |
| CTRL10 | Saline | 12.72 | 131.33 | 132 | 298.6 | Female |

This table shows the inoculum, maternal age at delivery, gestational age at inoculation and delivery, fetal body weight, and fetal sex for each animal in the study.

**Figure S1.** Proportion of Single-Cell Populations by Type of Control Group

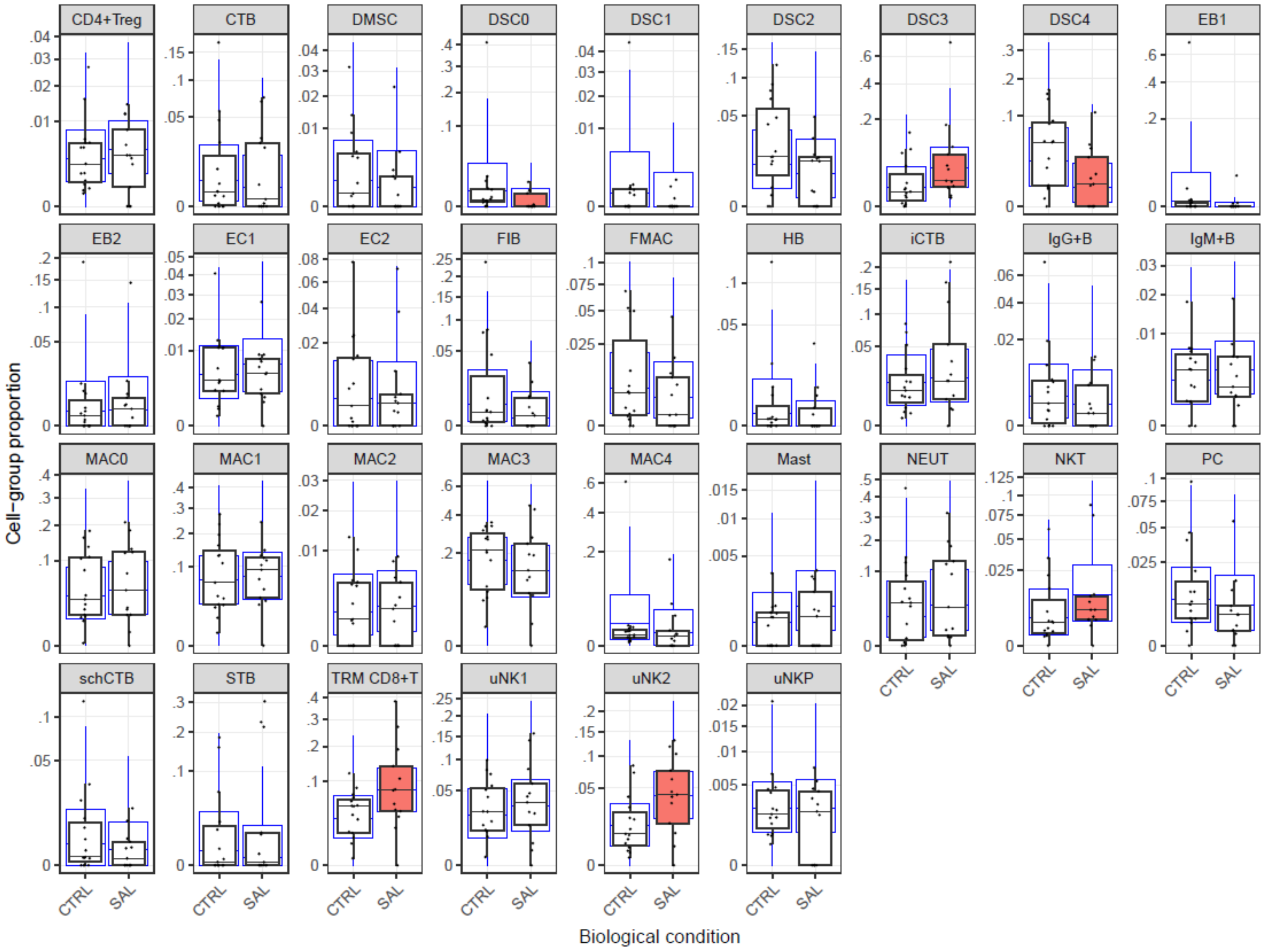


This figure illustrates the proportion of single-cell populations (y-axis) out of all single-cell populations for each type of control: subcutaneous media inoculation (CTRL) or choriodecidual saline (SAL) inoculation. Each dot within a bar plot represents a single sample from either the placental basal plate, chorioamniotic membranes or maternal decidua. The black box shows the median and the interquartile range. The blue bounding box indicates the mean +/- one standard deviation. A red colored box indicates a significant difference in proportions between the groups using a Fisher's exact test ($p<0.05$). Abbreviations for single-cell populations are shown in Figure 3.

**Figure S2.** Proportion of Single-Cell Populations by Tissue of Origin

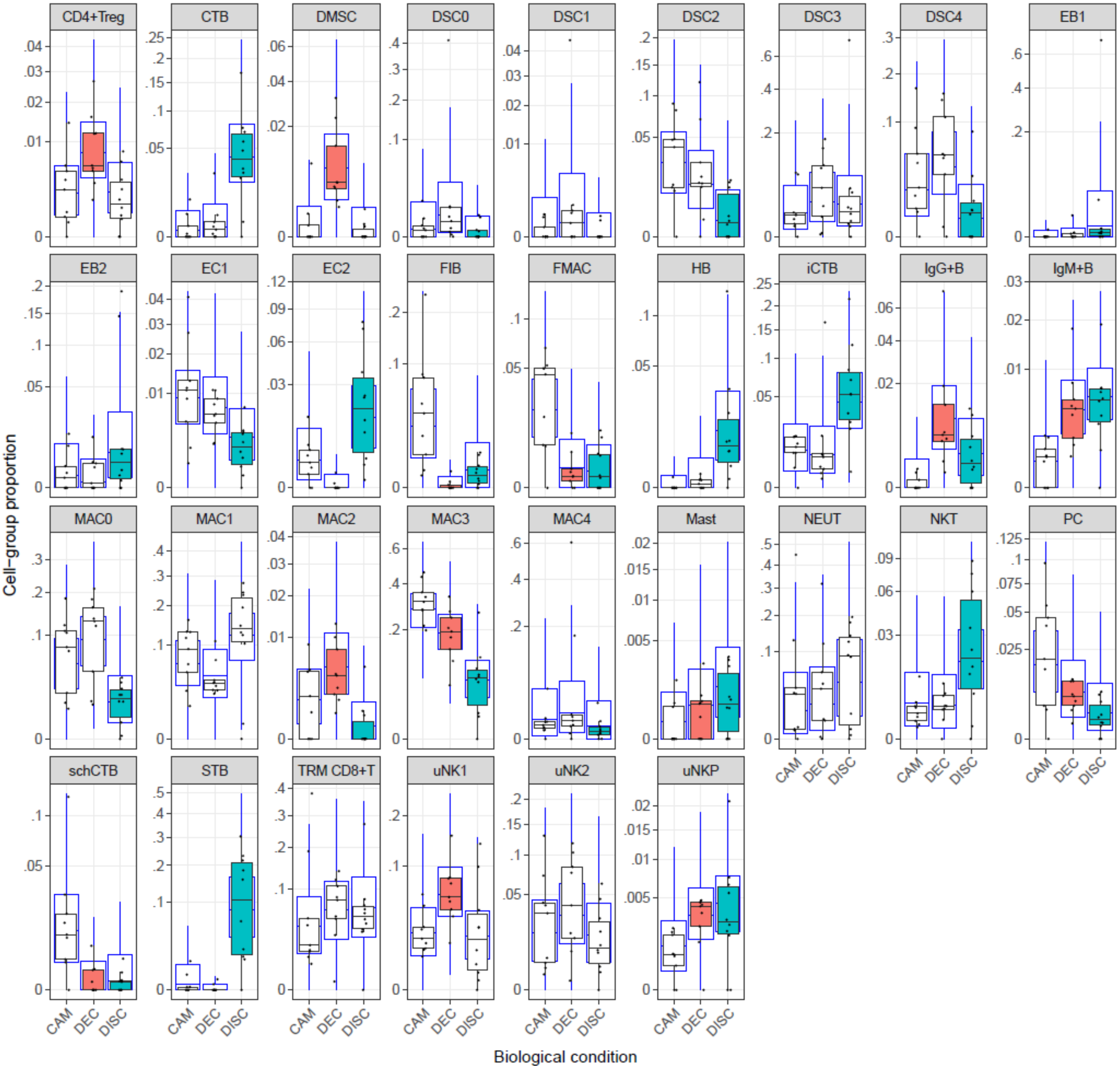


This bar plot illustrates the proportion of single-cell populations (y-axis) among all single-cell populations from each tissue source: placental disc (DISC), maternal decidua (DEC), and chorioamniotic membranes (CAM). Each dot within the bar plot represents a single sample from either the media or saline control groups. The black box shows the median and the interquartile range. The blue bounding box indicates the mean +/- one standard deviation. A colored box indicates a significant difference in proportions between either: 1) the DEC (red) and the CAM, or 2), the DISC (turquoise) and the CAM, using a constrained beta-binomial distribution ($p<0.05$). Abbreviations for single-cell populations are shown in Figure 3.

**Figure S3.** Proportion of Single-Cell Populations by Fetal Sex

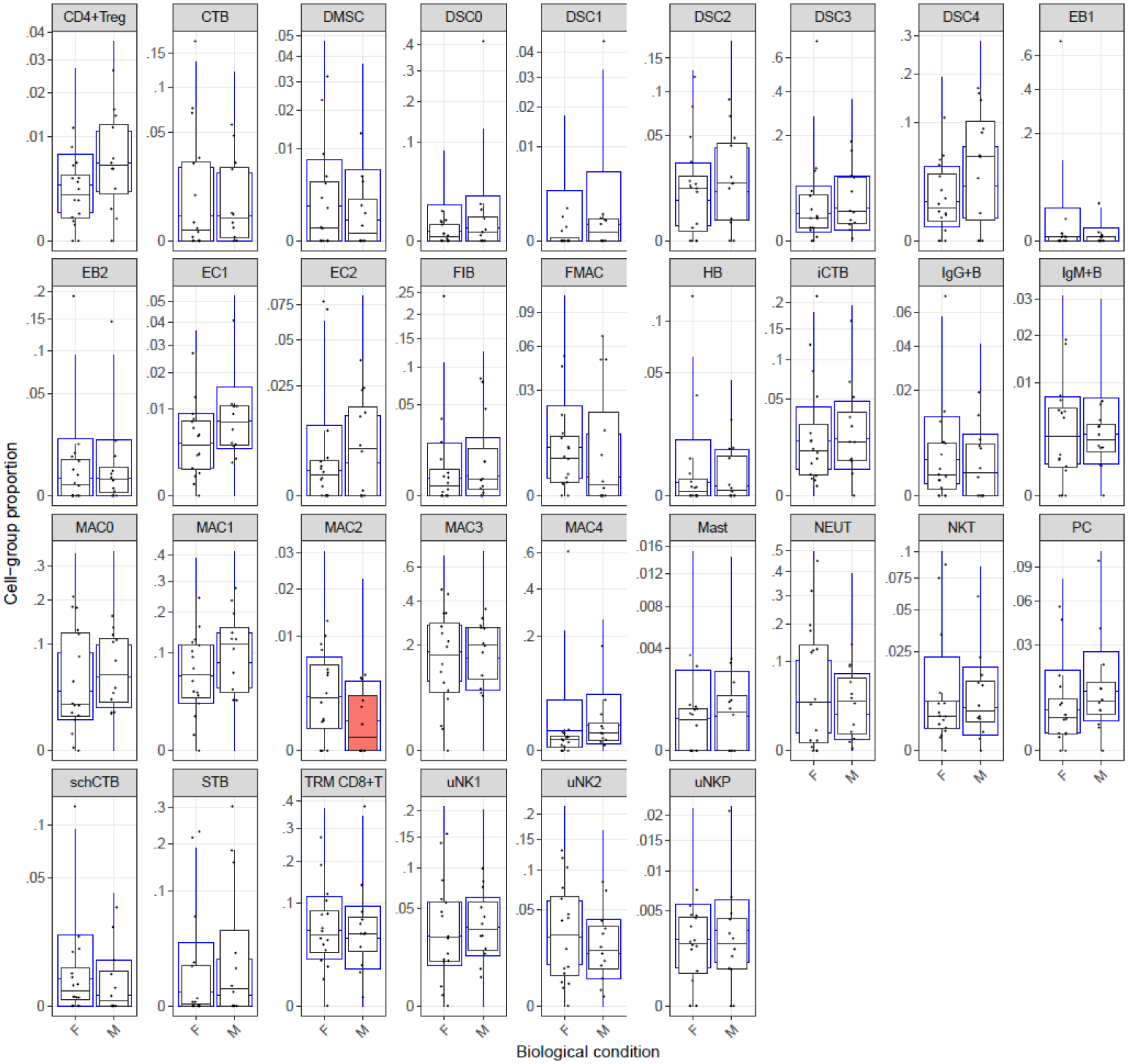


This bar plot illustrates the proportion of single-cell populations (y-axis) among all single-cell populations from each placenta group from a single sex: female (F) versus male (M). Each dot within the bar plot represents a single sample from either the media or saline control groups from a specific tissue (basal plate, membranes, decidua) grouped by fetal sex. The black box shows the median and the interquartile range. The blue bounding box indicates the mean +/- one standard deviation. A red colored box indicates a significant difference in proportions between placentas from a female versus a male fetus using a Fisher's exact test ($p<0.05$). Abbreviations for single-cell populations are shown in Figure 3.
